# Metabolic Dysfunction Underlying Fatigue in Multiple Sclerosis: Elevated Lactate and Impaired Post-Exercise Creatine Response in the Anterior Cingulate Cortex

**DOI:** 10.1101/2025.11.06.686949

**Authors:** Gemma Brownbill, Jeanne Dekerle, Mara Cercignani, Itamar Ronen, James Stone

## Abstract

**Background:** Multiple Sclerosis (MS) is characterised by altered brain metabolism and increased chronic perceived fatigue. However, the relationship between brain metabolites, fatigue, and physical task effects in MS remains unclear. This study investigated brain metabolite concentrations (glutamate + glutamine (Glx), lactate, and total creatine (tCr)) in the anterior cingulate cortex (ACC), a region involved in interoceptive processing, before and after a physical task.

**Methods:** Twenty-two people with MS (pwMS) and 22 controls underwent Magnetic Resonance Spectroscopy before and after fatiguing wrist extension tasks. Linear mixed models analysed group differences and exercise-induced metabolite changes.

**Results:** PwMS showed higher ACC lactate concentrations than controls at rest and post-exercise (*F* = 7.08, *p* = 0.011, 95% CI[0.033, 0.228]). No significant Glx differences were observed. A significant group × exercise interaction for tCr occurred (*F* = 4.63, *p* = 0.037, 95% CI[0.027, 0.581]), with tCr decreasing post-exercise in controls (*t* = 3.09, *p* = 0.02) but remaining stable in pwMS. Changes in lactate correlated moderately with perceived effort in pwMS (*r* = 0.51, *p* = 0.04).

**Conclusions:** This study provides novel evidence of metabolic differences in pwMS, characterised by elevated lactate and stable post-exercise tCr, suggesting altered energy metabolism potentially linked to mitochondrial dysfunction.

## Introduction

Fatigue is one of the most disabling symptoms in multiple sclerosis (MS), affecting quality of life independently of physical disability (1,2). This chronic, pathological fatigue, often referred to as trait fatigue, represents the overall disposition and intensity of perceived fatigue experienced over extended periods and is typically unalleviated by rest (3). Despite its prominence, the underlying neurobiological mechanisms of MS-related fatigue remain poorly understood, with no clear consensus on the neurological changes that lead to fatigue (2).

Recent research has highlighted the potential role of altered brain metabolites in MS-related trait fatigue, particularly involving energy substrates and neurotransmitter systems (3). Importantly, fatigue, defined as a “feeling of diminishing capacity to cope with physical or mental stressors, either imagined or real” (4), encompasses both a chronic component (trait fatigue) and a ‘dynamic component’ (state fatigue), the latter representing momentary sensations that can change rapidly. To investigate neurobiological mechanisms underlying both trait and state fatigue, physical exercise paradigms can acutely induce state fatigue in individuals with pre-existing chronic fatigue. This allows examination of brain metabolite changes, measured by Proton Magnetic Resonance Spectroscopy (1H-MRS), in response to fatiguing tasks, potentially revealing mechanisms underlying MS-related fatigue and responses to acute challenges.

Among the brain metabolites implicated in MS, glutamate and those involved in energy metabolism play a particularly significant role in disease progression and symptom manifestation (5–9). Glutamate excitotoxicity is reported as one of the pathological mechanisms of MS, resulting in oligodendrocyte and glial cell death (5). Brain glutamate + glutamine concentrations (Glx) have been associated with disability in MS, with higher levels in the sensorimotor and parietal region correlating with greater physical and cognitive impairment (10,11). However, despite this evidence linking metabolites to MS pathology, only one study has directly linked brain metabolites to perceived fatigue in MS, showing higher Glx and lower γ-aminobutyric acid (GABA) concentrations in prefrontal and sensorimotor cortices of pwMS with elevated trait fatigue scores (3).

In parallel, mitochondrial dysfunction and altered energy metabolism are increasingly recognised as contributors to MS pathology (6). Increased glycolytic activity results in elevated cerebrospinal fluid (CSF) lactate concentrations associated with disease progression (7,8). Creatine and phosphocreatine metabolism are also altered in MS, with studies reporting both elevated and decreased concentrations in various brain regions depending on disease stage and lesion location (9). However, the relationship between brain energy metabolism and fatigue perception remains unexplored in MS, and it is unclear whether baseline metabolic alterations translate to differential responses during acute metabolic challenges, such as exercise-induced fatigue.

1H-MRS has been used to study exercise-induced brain metabolite changes in healthy populations. A recent systematic review identified nine studies examining these effects, with exercise ranging from yoga to vigorous cycling (12). Results show tendencies for GABA and lactate concentrations to increase post-exercise.

Vigorous cycling increases Glx in the visual cortex of healthy controls (13,14). This may reflect heightened neuronal activity as glutamate serves as the primary excitatory neurotransmitter (15). However, small sample sizes and inconsistent methodologies limit generalisability, and evidence remains inconclusive for glutamate and creatine which, as noted above, are relevant to MS pathology. Furthermore, these studies have focused primarily on metabolite changes without examining concurrent changes in perceived state fatigue, limiting understanding of how biochemical and subjective responses relate.

Additionally, two studies confirmed post-exercise brain lactate increases in the visual and occipital cortex after vigorous cycling (14,16), as lactate can be utilised by the brain as an energy substrate during exercise (17). Total creatine (tCr) is also an important brain energy substrate (18) that is normally assumed to be a stable metabolite in 1H-MRS studies. Interestingly, one study found decreased tCr in the occipital cortex (16) after vigorous cycling; however, another found no change with similar exercise (14). Studies examining exercise-induced neurometabolic changes are lacking in pwMS, despite the potential to uncover possible disease-related fatigue mechanisms.

One brain region of particular interest for the study of state and trait fatigue is the anterior cingulate cortex (ACC), which plays a key role in interoception (the perception of internal bodily signals) and likely, fatigue perception (19,20). The ACC receives inputs from the insula and plays a crucial role in performance monitoring during physical challenges (21). During exercise-induced fatigue, the ACC integrates interoceptive signals that likely inform state fatigue and effort perception (defined as “the conscious sensation of how hard, heavy and strenuous exercise is” (22)), helping maintain homeostasis (20,22,23). The dorsal ACC (dACC) is particularly relevant for studying perceived fatigue due to its involvement in interoceptive (23) and metacognitive pathways (19), and may show altered metabolic activity in MS-related trait and state fatigue (24).

Despite existing (but scarce) research on brain metabolite differences in pwMS and exercise-induced changes in healthy individuals, the relationship with both trait and state fatigue remains unclear. This study aims to investigate neurometabolic changes in the dACC of pwMS vs healthy controls before and after fatiguing isometric wrist extension contractions with a focus on Glx, lactate, and tCr concentrations. Based on previous evidence, it was hypothesised that pwMS would exhibit higher baseline concentrations of Glx and lactate in the ACC, and that metabolite responses to a fatiguing physical task would differ between pwMS and healthy controls.

## Methods

### Participants

Ethical approval was granted by the Health and Social Care Research Ethics Committee B (Northern Ireland) (REC reference: 19/NI/0154). Twenty-two relapsing-remitting pwMS (3 male), and 22 age and sex-matched controls were recruited and gave written informed consent. Patient characteristics can be viewed in Table 1. MS patients had an EDSS < 3.5 and were > 2 months clear from their most recent relapse. All participants had no contraindications to MRI, and control participants had no history of cardiovascular, neurological or musculoskeletal disorders.

**Table 1.** Multiple sclerosis and healthy control participant demographics.

|  | Multiple Sclerosis Group<br>(*n* =22) | Control Group<br>(*n* =22) |
| --- | --- | --- |
| Age (years), mean ± SD | 47.7 ± 8.9 | 48.4 ± 9.5 |
| Age range (years) | 30–60 | 29–58 |
| Sex, *n* | 3 males, 19 females | 3 males, 19 females |
| MS disease duration (years), mean ± SD (range) | 9.8 ± 8 (0.67–31) | N/A |
| Disease modifying therapy, *n* | 9 none; 3 Ocrelizumab; 2 Tysabri; 2 Aubagio; 1 Kesimpta; 1 Fingolimod | N/A |
| FSS score (average ± SD) | 4.99 ± 0.95* | 2.71 ± 0.88 |
| Baseline RoF (average ± SD) | 2.72 ± 0.98* | 1.68 ± 1.67 |

MS disease duration was calculated from diagnosis. FSS, Fatigue Severity Scale; RoF, Rating of Fatigue; SD, standard deviation; MS, multiple sclerosis;
N/A, not applicable.\* significantly different (*p*<0.05)

### Experimental design

Participants performed two bouts of sustained isometric wrist extension contractions inside the MRI scanner: A 3-minute all-out contraction (3AO) was followed by a 5-minute submaximal contraction (5SUB) at 55% of the average force maintained over the last 30s of the 3AO; the two bouts of exercise were separated by 8 minutes of rest. 1H-MRS acquisition runs were performed before the 3AO and following the 5SUB.

### MRI-compatible ergometer

A custom 3D-printed MRI-compatible ergometer secured the forearm. The wrist was secured with straps, and an MRI force dynamometer (MP3X; Biopac, CA, USA) was positioned 1 cm below the middle metacarpophalangeal joint. The arm was positioned parallel to the body with the elbow extended.

### Perceptual measures

The Fatigue Severity Scale (FSS) (1) was used to measure trait perceived fatigue upon arrival to the laboratory. The Rating of Fatigue scale (RoF) (4) was then used to measure state perceived fatigue before and after the physical task. The CR-100 scale (25) was used to measure rating of perceived effort (RPE) every 30 s during 5SUB. The scale was presented on screen, and participants verbally reported their RPE. Participants were instructed on the definitions of fatigue (4) and effort (22), emphasising the distinction between effort (instantaneous exertion intensity) and fatigue (overall capacity to continue).

### MRI and 1H-MRS acquisition

1H-MRS scans were performed on a 3T Siemens Magnetom Prisma scanner (Siemens Healthineers, Erlangen, Germany) using a 32-channel head coil. Participants were positioned supine in the scanner with their arm secured in the custom-built wrist extensor ergometer. After a standard localiser sequence, a T1-weighted magnetisation prepared rapid gradient-echo (MPRAGE) structural scan was acquired (TR=2,300ms, TE=2.19ms, 1mm isotropic resolution, 192 sagittal slices).

1H-MRS data were acquired using a plug-and-play automatic semi-localisation by adiabatic selective refocusing (sLASER) sequence (26). The main acquisition parameters included a TE of 30 ms and a TR of 3000 ms, with a spectral bandwidth of 2500 Hz. A total of 64 averages were collected, along with four non-water-suppressed acquisitions. The voxel of interest (VOI) was positioned on the dACC, with a size of approximately 9 cc (30 × 15 × 20 mm³). B_0_ shimming was performed using FASTESTMAP, ensuring optimal magnetic field homogeneity. Water suppression was achieved with VAPOR (Variable Power and Optimised Relaxation Delays) (27), with B_0_ shimming and RF pulse calibrations performed automatically.

### Data processing and analysis

Spectra were processed using LCModel with linear combination modelling. Pre-processing steps included eddy current correction using the non-water-suppressed data and spectral registration across averages to correct for frequency and phase drifts (28). A basis set simulated for sLASER acquisitions at 3T with TE = 30ms was used for spectral fitting, including major brain metabolites, macromolecules, and lipids. Bespoke Matlab (Mathworks, Natick, MA) scripts generated LCModel-compatible RAW files for analysis. Data were scaled to water and corrected for tissue-specific T1 and proton density values for grey matter, white matter, and CSF using values from (29). Tissue segmentation was performed on the MPRAGE volume using SPM12 via Gannet, generating VOI-specific tissue fractions that were incorporated into the LCModel water scaling factor (Figure 1; B & C). Metabolite estimates with Cramer-Rao Lower Bounds > 20% were excluded, and spectra were visually inspected for quality (Figure 1; A).

**Figure 1.**
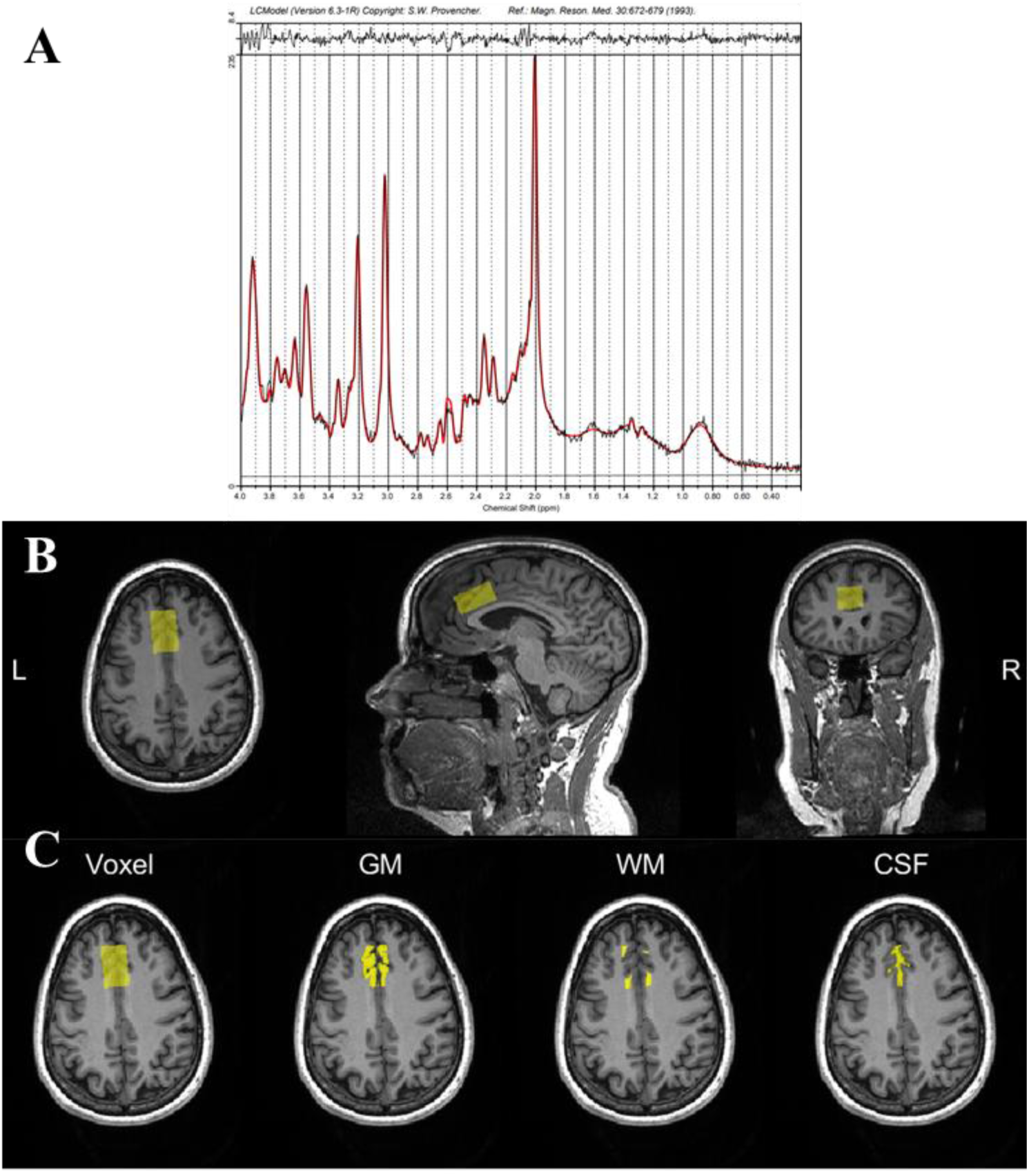
1H-MRS Gannet post-processing output and example spectra. A: example spectra with the fit line. The red line shows the fitted data, and the black line the raw data. B: 1H-MRS voxel positioning on the coregisted T1 weighted image for an example subject. C: Gannet segment of the segmented different tissue types within the voxel of interest on an axial T1 weighted image. GM: grey matter. WM: white matter. CSF: cerebrospinal fluid.

### Statistical analysis

Data normality was assessed using Shapiro-Wilk tests and Q-Q plots, with outliers identified via standardised residuals and influence diagnostics. Linear mixed models (LMMs) were performed using Jamovi (GAMLj module) with participant as a random factor and group (pwMS vs. controls) and time (pre-vs. post-exercise) as fixed factors. Effect sizes were reported as partial eta-squared (ηp²) with Satterthwaite approximation for degrees of freedom. Pearson correlations examined associations between metabolite concentrations (baseline and post-exercise) and fatigue measures (FSS and RoF), and between metabolite percentage changes and changes in state fatigue (RoF) and perceived effort.

## Results

One MS patient was excluded due to poor spectral quality, and several recordings in pwMS were removed due to high Cramer-Rao Lower bounds (5 lactate, 2 Glx, 2 tCr recordings; Supplementary Table S1). In the control group, only one lactate recording was removed post-exercise. Averages and SD for metabolites of interest can be viewed in Table 2, with complete metabolite data presented in Supplementary Table S2.

**Table 2.** Average metabolite concentrations (± standard error of the mean) for glutamate + glutamine, lactate and total creatine.

|  |  | Pre-exercise | Post-exercise |
| --- | --- | --- | --- |
| | | Metabolite concentration<br>(± SE) | Metabolite concentration<br>(± SE) |
| Glx (mmol/kg) | CG | 13.4 (0.17) | 13.1 (0.19) |
|  | MS | 13.8 (0.37) | 13.6 (0.51) |
| Lactate (mmol/kg) | CG | 0.75 (0.035) | 0.69 (0.036) |
|  | MS | 0.97 (0.048) * | 0.91 (0.046) * |
| tCr (mmol/kg) | CG | 9.34 (0.10) † | 9.04 (0.14) † |
|  | MS | 9.30 (0.10) | 9.23 (0.18) |

Glx: glutamate + glutamine; tCr: total creatine; SE: standard error; MS: multiple sclerosis; CG: control group; \*: significant effect of group; †: significant group\*time interaction.

PwMS demonstrated significantly higher trait (FSS: 4.99 ± 0.95 vs. 2.71 ± 0.88, *t*₍_42_₎ = -8.21, *p* < 0.001, *d* = -2.45) and baseline state fatigue (RoF: 2.72 ± 0.98 vs. 1.68 ± 1.67, *t*₍₃₄.₅₎ = - 2.35, *p* = 0.02, *d* = -0.70) compared to controls. State fatigue increased significantly post-exercise (pwMS: 6.0 ± 1.87; controls: 3.59 ± 1.96; *F*₍₅,₂₀₉₎ = 43.86, *p* < 0.001, ηₚ² = 0.51), with persistently higher levels in pwMS (*F*₍₁,₄₂₎ = 16.28, *p* < 0.001, ηₚ² = 0.28) and a significant group × time interaction (*F*₍₅,₂₀₉₎ = 3.12, *p* = 0.010, ηₚ² = 0.07), indicating greater increases in pwMS. RPE increased significantly over time (*F*₍₉,₃₉₆₎ = 90.1, *p* < 0.001, ηₚ² = 0.67) with no group effect (*F*₍₁,₄₃.₅₎ = 2.51, *p* = 0.12) but a significant group × time interaction (*F*₍₉,₃₉₆₎ = 3.47, *p* < 0.001, ηₚ² = 0.07).

There was no significant effect of group for Glx (*F*_(1, 39.8)_ = 0.82, *p* = 0.360, SE = 0.31, 95% CI [-0.333, 0.911], η_p_^2^ = 0.02). The effect of time was marginally not significant (*F*_(1, 39.5)_ = 2.98, *p* = 0.092, SE = 0.230, 95% CI [-0.849, 0.054], η_p_^2^ = 0.07). There was no significant interaction effect between group and time (*F*_(1, 39.5)_ = 0.128, *p* = 0.722, SE = 0.46, 95% CI [-1.067, 0.737], η_p_^2^ = 0.003). Figure 2 depicts Glx pre- and post-exercise in individuals in both groups.

**Figure 2.**
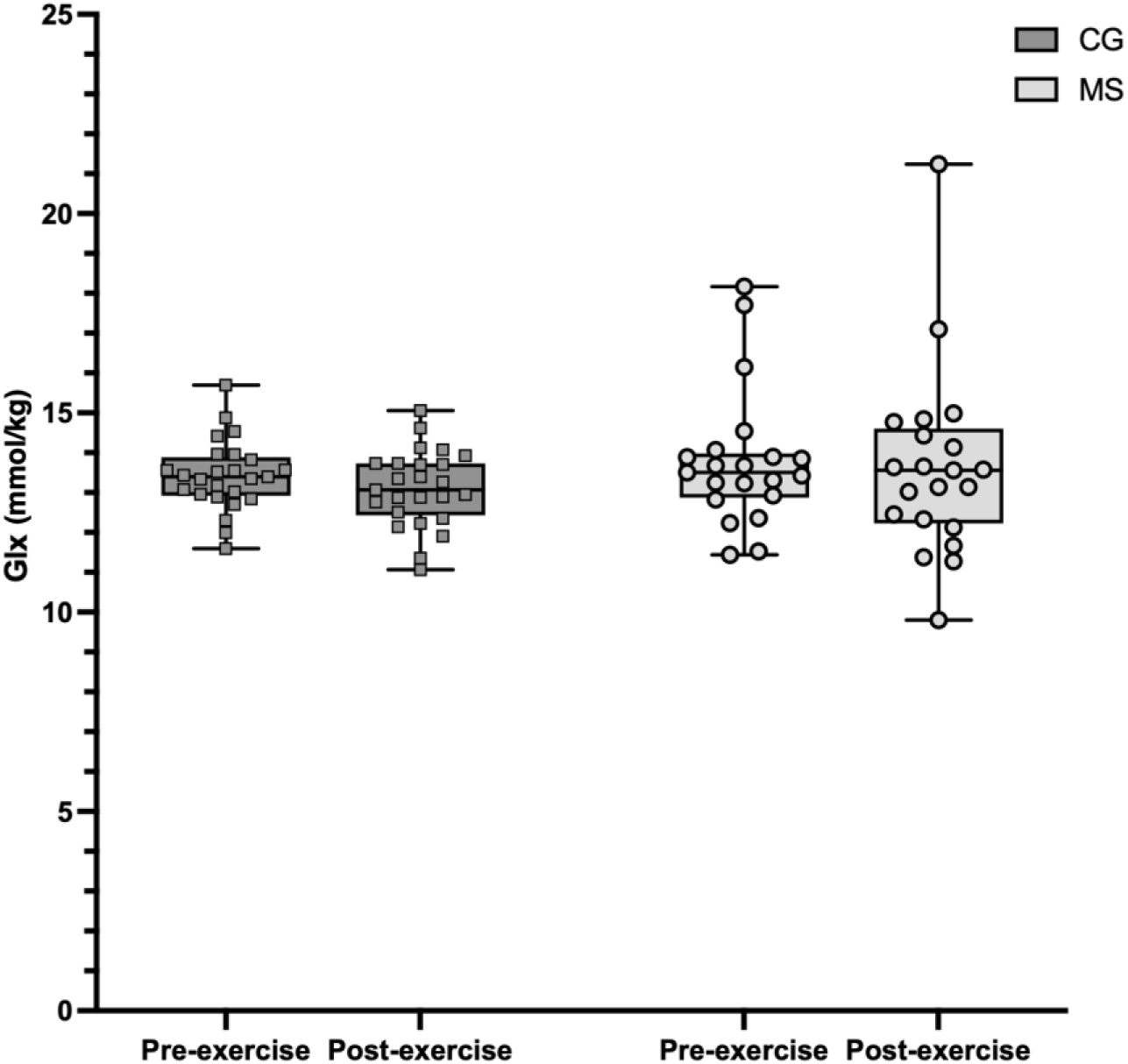
Comparison of glutamate + glutamine concentrations in control and multiple sclerosis groups before and after exercise. Glx: glutamate + glutamine; CG: control group; MS: multiple sclerosis.

There was a significant effect of group for lactate (*F*_(1, 42.3)_ = 7.08, *p =* 0.011, SE = 0.04, 95% CI [0.033, 0.228], η ^2^ = 0.14). There were no significant effects of time (*F*_(1, 40.9)_ = 1.74, *p =* 0.194, SE = 0.03, 95% CI [-0.093, 0.019], η ^2^ = 0.04) or interaction between group and time (*F*_(1, 40.9)_ = 0.13, *p =* 0.72, SE = 0.60, 95% CI [-0.096, 0.140], η_p_^2^ = 0.003). Individual values are presented in Figure 3.

**Figure 3.**
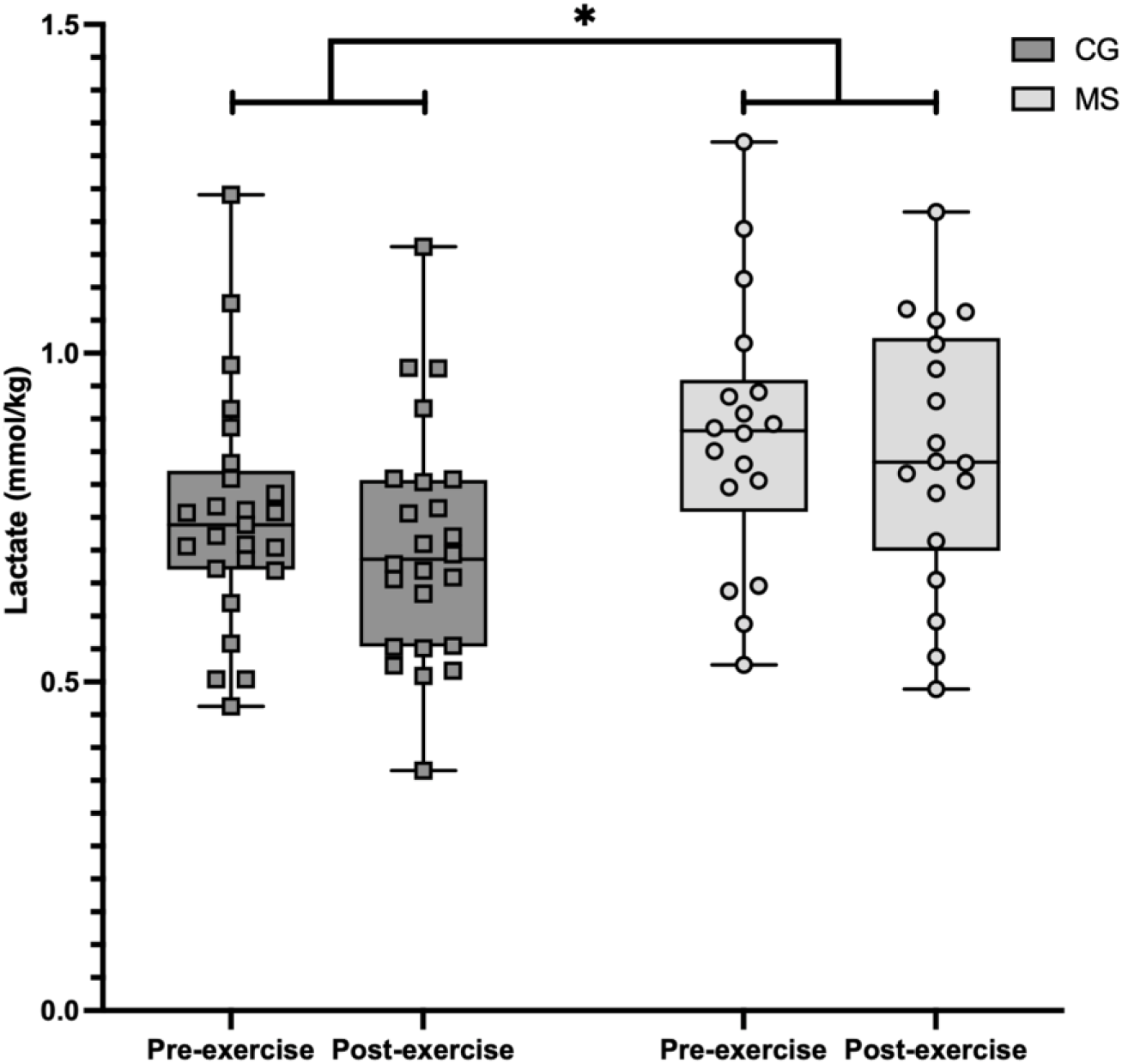
Comparison of lactate concentrations in control and multiple sclerosis groups before and after exercise. CG: control group; MS: multiple sclerosis; *: *p* < 0.05.

For tCr, there was no effect of group (*F*_(1, 42.1)_ *=* 0.03, *p* = 0.849, SE = 0.18, 95% CI [-0.332, 0.404], η ^2^ = 0.0007). The effect of time was marginally not significant (*F*_(1, 41.3)_ *=* 3.94, *p* = 0.054, SE = 0.07, 95% CI [-0.279, -0.001], η ^2^ = 0.08). However, there was a significant group x time interaction effect (*F*_(1, 41.3)_ *=* 4.63, *p* = 0.037, SE = 0.14, 95% CI [0.027, 0.581], η_p_^2^ = 0.10). Post-hoc analysis revealed that tCr significantly reduced post exercise in the CG only (*t*(_43_) = 3.09, *p* = 0.02, *d* = 0.47, SE = 0.09). The decline in the CG can be viewed in Figure 4.

**Figure 4.**
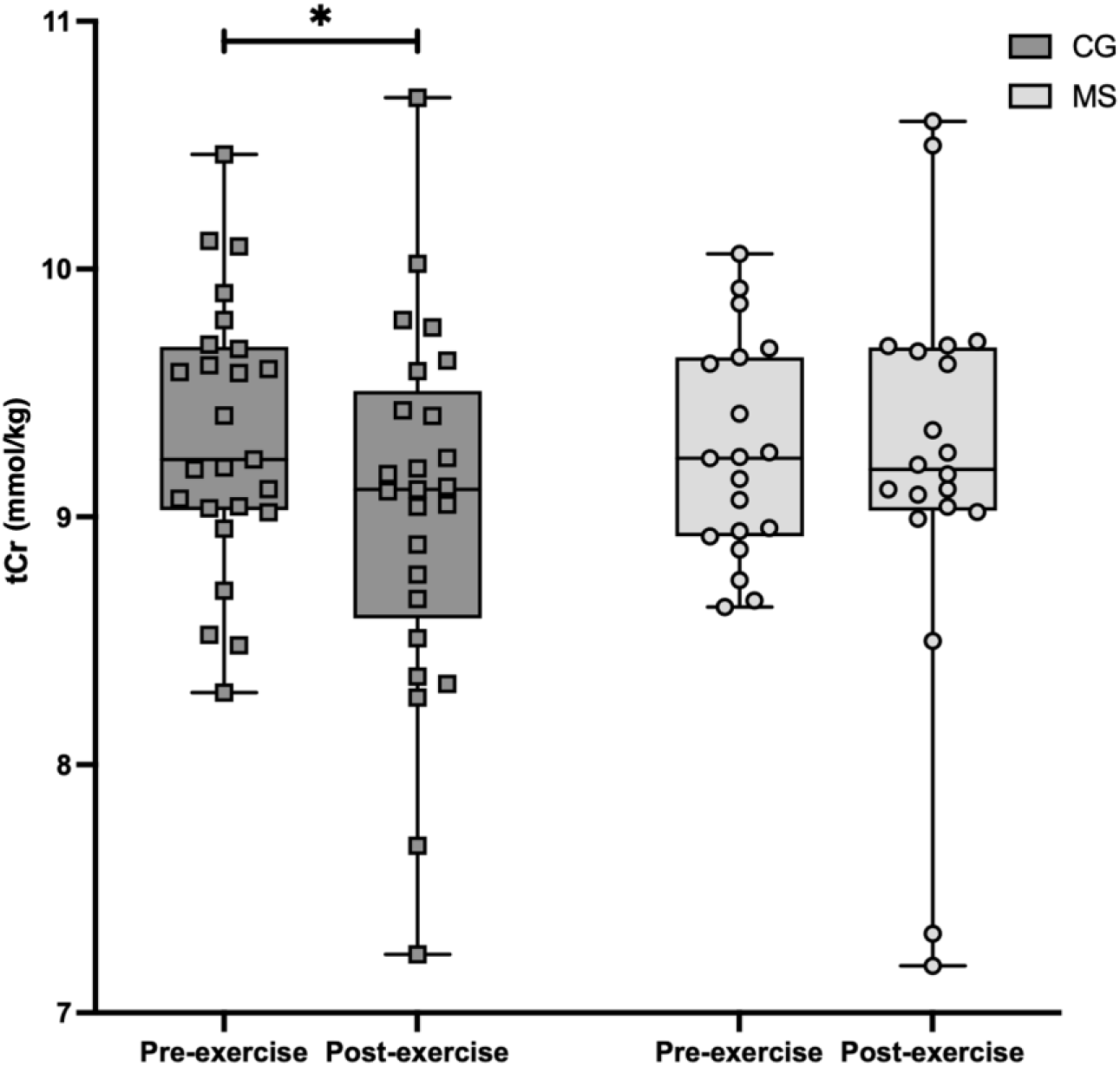
Comparison of total creatine concentrations in control and multiple sclerosis groups before and after exercise. tCr: total creatine; CG: control group; MS: multiple sclerosis; *: *p* < 0.05.

No significant correlations were found between baseline metabolite concentrations and state (RoF) or trait (FSS) fatigue measures. Similarly, post-exercise metabolite concentrations showed no associations with post-exercise RoF in either group. However, a moderate positive correlation was observed between lactate changes and perceived effort in pwMS only (Figure 5; *r* = 0.51, *p* = 0.04).

**Figure 5.**
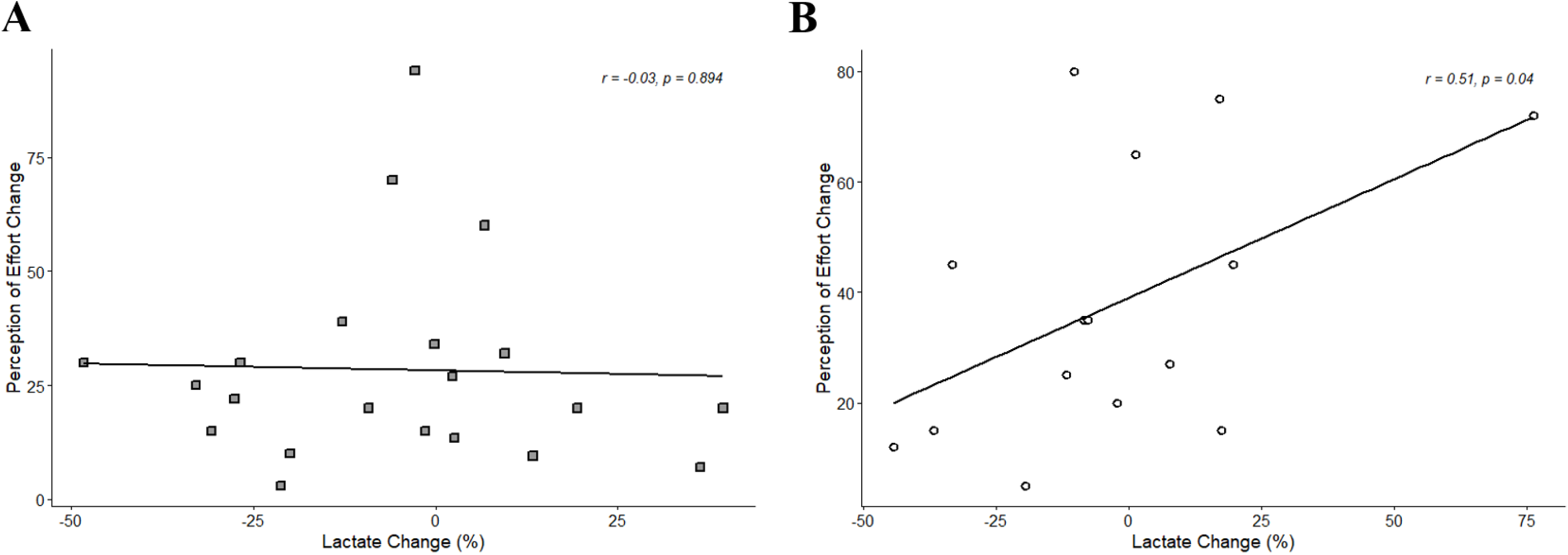
Correlation between lactate and perception of effort changes during fatiguing exercise. Relationship between percentage change in lactate concentrations and perception of effort in (A) healthy controls (*n* = 21) and (B) people with multiple sclerosis (*n* = 18). Each point represents an individual participant. Pearson correlation coefficients (*r*) and *p*-values are shown in each panel. A significant positive correlation was observed only in the multiple sclerosis group.

## Discussion

This study provides the first evidence of metabolic changes in the ACC of pwMS following fatiguing wrist extensions, revealing chronically elevated lactate concentrations (at both baseline and post-exercise) and differential tCr exercise responses compared to healthy controls, while showing no group differences in Glx concentrations. These findings suggest metabolic dysfunction in the ACC of pwMS, offering novel insights into the metabolic underpinnings of MS-related fatigue.

PwMS demonstrated significantly higher trait and baseline state perceived fatigue compared to controls, consistent with established literature (1,2). Both groups showed increased state fatigue and effort perception following the exercise protocol, with pwMS exhibiting persistently higher levels and greater increases compared to controls (post exercise RoF; CG: 3.59 ± 1.96; pwMS: 6.0 ± 1.87), indicating the exercise protocol induced a greater perceived fatigue response in pwMS. Importantly, baseline metabolite concentrations did not correlate with trait or state fatigue measures in either group, indicating that while metabolic alterations exist in the ACC of pwMS, they may not directly drive perceived fatigue sensations at rest.

Similar ACC Glx levels in pwMS and controls, both at baseline and post-exercise, contrast with elevated glutamate reported in other MS brain regions (10,11,30). Given the ACC’s essential role in interoceptive awareness, stable glutamate concentrations may represent a compensatory mechanism to maintain critical functions despite ongoing disease processes. The low disability levels in our MS sample (EDSS <3.5) may also account for preserved glutamatergic function in this region.

The significantly elevated lactate concentrations in pwMS at rest and following physical exercise represent the first in vivo evidence of higher brain lactate in the ACC in this population. This finding aligns with studies showing elevated CSF lactate in MS (7,8), and may indicate ongoing neuroinflammatory processes in this critical brain region involved in interoceptive processing and fatigue perception in pwMS (24). Importantly, these elevated lactate levels did not increase post-exercise and showed no associations with fatigue measures at any time point, suggesting that while lactate reflects underlying metabolic dysfunction, it may not directly influence fatigue perception. This disconnect between metabolic markers and subjective fatigue suggests that MS-related fatigue arises from multifaceted mechanisms beyond metabolic alterations alone.

The absence of post-exercise lactate increases contrasts with previous whole-body exercise studies (14,16). The single-joint contraction protocol generated moderate metabolic demands sufficient to induce perceived fatigue while involving lower muscle mass recruitment and ATP requirements than whole-body exercise, resulting in correspondingly reduced blood lactate increases and brain lactate uptake (17,31).

In contrast to the stable lactate and Glx responses after exercise, whilst tCr was similar pre-exercise, it showed a significant differential pattern between groups. Controls demonstrated decreased post-exercise tCr, likely reflecting efficient phosphocreatine utilisation as an energy source (18). This finding is consistent with Dennis *et al*.’*s* (16) finding in the occipital cortex, and challenges the traditional view of tCr as a stable internal reference and supports absolute quantification methods in metabolic studies. Conversely, pwMS showed stable tCr levels post-exercise, suggesting dysregulated creatine metabolism (9) or potential allocation of tCr for neuroprotective processes rather than energy production (32). This inability to mobilise tCr during metabolic challenges may reflect dysregulated energy metabolism in MS.

The combined findings of elevated lactate concentrations and differential tCr exercise responses in pwMS indicate underlying mitochondrial dysfunction, a well-documented feature of MS pathophysiology (6,33). Mitochondrial impairment leads to reduced oxidative phosphorylation efficiency, increasing reliance on glycolysis and resulting in elevated lactate production. Combined with the stable post-exercise tCr discussed above, this metabolic inefficiency in the ACC, a region critical for interoceptive processing and fatigue perception (20,24), may contribute to the elevated baseline fatigue and greater exercise-induced fatigue increases observed in pwMS.

Effort perception increased at a greater rate and was correlated with lactate changes in pwMS only (*r* = 0.51, *p* = 0.04). These observations could be interpreted through the ACC’s dual role in interoceptive processing and effort perception (19,22,23). Metabolic dysfunction in this region, evidenced by chronically elevated lactate and impaired creatine utilisation, may disrupt sensory signal processing. Lactate modulates neuronal excitability by reducing cortical calcium spiking frequency (34), potentially affecting the brain’s capacity to interpret internal bodily signals during exertion. This disrupted interoceptive processing may create a feedback loop where heightened effort perception reinforces fatigue through impaired sensory attenuation (the brain’s reduced ability to suppress self-generated sensory feedback) and altered metacognitive evaluation of homeostatic control (19,20,35). Metabolic efficiency variations may modulate effort through these pathways.

This study has several limitations. The modest sample size limits statistical power and restricts generalisability to low-disability relapsing-remitting MS. Methodologically, the approximately 4-minute delay between exercise cessation and MRS data collection, necessitated by scanner calibration procedures, may have allowed partial metabolite normalisation. This may account for the absence of post-exercise lactate increases in both groups, contrasting with previous whole-body exercise studies. Lastly, the focus on a single brain region may not capture whole-brain responses.

This study provides novel evidence of metabolic differences in the ACC of pwMS, characterised by elevated lactate levels both at rest and following a physical exercise, and impaired tCr utilisation post-exercise. While these metabolic alterations do not directly correlate with perceived fatigue and effort, they suggest underlying mitochondrial dysfunction that may contribute to the complex pathophysiology of MS-related fatigue. The differential metabolic responses to exercise highlight the potential for targeted interventions addressing energy metabolism in MS fatigue management.

## Statements and declarations

The authors declared no potential conflicts of interest with respect to the research, authorship, and/or publication of this article.

## Author Contributions

G.B., M.C. and J.D. conceptualised the study. G.B. conducted data collection and analysis and wrote the manuscript. J.D., M.C., and J.S. supervised the research, providing guidance on study design, methodology, and interpretation. I.R. provided MRS technical expertise and contributed to data processing protocols. All authors contributed to manuscript revisions and approved the final version.

## Data availability statement

The raw data supporting the conclusions of this article will be made available by the authors on request.

## Acknowledgements

We would like to thank Dr Andrew Barritt for assisting with participant recruitment, and the laboratory and imaging technicians at the University of Brighton and Brighton and Sussex Medical School for their support during conceptualisation and data collection, and the volunteers who participated. We thank Dr. Malgorzata Marjanska and Dr. Edward Auerbach from the Center for Magnetic Resonance Research (CMRR) at the University of Minnesota for providing us with the CMRR MRS sequence package, and Dr. Dinesh Deelchand from the CMRR for providing us with the plug-and-play sLASER sequence and the LCModel basis set.

