## Supplemental Table 1 for "Metabolic Dysfunction Underlying Fatigue in Multiple Sclerosis: Elevated Lactate and Impaired Post-Exercise Creatine Response in the Anterior Cingulate Cortex"

S1: Average (± SD) Cramer-Rao Lower Bounds for glutamate + glutamine, lactate and phosphocreatine + creatine.

| | Pre-exercise | | Post-exercise | |
|---|---|---|---|---|
| Metabolite | MS | CG | MS | CG |
| Glx (%) | 2.32 (± 0.47) *n* = 20 | 2.08 (± 0.27) *n* = 22 | 2.45 (± 0.67) *n* = 20 | 2.08 (± 0.27) *n* = 22 |
| Lactate (%) | 17.1 (± 2.08) *n* = 18 | 17.0 (± 2.86) *n* = 22 | 17.6 (± 3.03) *n* = 18 | 18.3 (± 2.54) *n* = 21 |
| tCr (%) | 1 (± 0) *n* = 20 | 1 (± 0) *n* = 22 | 1 (± 0) *n* = 20 | 1 (± 0) *n* = 22 |

Glx: glutamate + glutamine; tCr: total creatine; MS: multiple sclerosis; CG: control group; *n:* number of participants included in the analysis.
